# MOAflow: how re-design a pipeline with Nextflow streamlines data analysis

**DOI:** 10.64898/2026.03.26.713914

**Authors:** Jacopo Tartaglia, Mario Giorgioni, Luigi Cattivelli, Primetta Faccioli

**Affiliations:** CREA - Research Centre for Genomics and Bioinformatics, Fiorenzuola d’Arda (PC), Italy; CREA - Research Centre for Forestry and Wood, Roma, Italy

**Keywords:** Nextflow, MOA-seq, high-performance computing, cloud computing, big data, workflow management systems

## Abstract

**Background:** Advances in high-throughput DNA sequencing technologies have dramatically reduced the time and cost required to generate genomic data. As sequencing is no longer a limiting factor, increasing attention must be paid to optimizing the analyses of the large-scale datasets produced. Efficient processing of such data is essential to reduce computational time and operational costs. In this context, workflow management systems (WMSs) have become key instruments for orchestrating complex bioinformatic pipelines. Among these systems, Nextflow has emerged as one of the most widely adopted solutions in bioinformatics.

**Methods:** To improve scalability and computational efficiency, we employed Nextflow to re-design an already existing pipeline dedicated to the analysis of MNase‐defined cistrome‐Occupancy (MOA‐seq) data. The re‐engineering process focused on modularizing the workflow and integrating containerization technologies to ensure reproducibility and easier deployment across heterogeneous computing environments.

**Results:** The resulting workflow, named MOAflow, represents a modernized and fully containerized pipeline for MOA‐seq data analysis. With only Docker and Nextflow required, the pipeline guarantees high portability and reproducibility. The data of the original article was used to benchmark the new pipeline. Its outputs closely match those of the original study with minor variations.

**Conclusions:** MOAflow demonstrates how the adoption of robust WMS can substantially enhance the performance and usability of pre-existing bioinformatic pipelines. By leveraging containerization and Nextflow, it ensures consistent results across platforms while minimizing setup complexity. This work highlights the value of modern WMS-driven approaches in meeting the computational demands.

## Introduction

The progressive reduction in costs and the easier access to high-throughput sequencing technologies have led to massive data production in biology and in its omics disciplines. Accordingly, nowadays the bottleneck lies in the bioinformatic analysis of data and no longer in their production (*Berger et al., 2023*). This situation has highlighted the need to increase the processing and storage capacity dedicated to data handling. However, meeting this challenge depends not only on implementing additional IT resources, but also on improving the efficiency with which they are used (*Berger et al., 2023*). In this framework workflow management systems play a central role in streamlining data analysis. Nextflow (*Di Tommaso et al., 2017*), which is a scientific workflow system based on the new DSL2 syntax, simplifies the writing of data-intensive computational pipelines. Nextflow has many bioinformatics applications, because it offers several advantages, including mature support for container technologies (e.g. Docker, Singularity…), compatibility with multiple source-code management systems, and a reactive programming model that enables highly efficient parallel execution and scalable workflows. Each process is organized as an independent module, promoting clarity and code reusability, while also allowing for easy implementation. The various processes are linked together in a workflow script, which can also be divided into sub-workflows, giving the code a transparent design that makes Nextflow pipelines easy to read, modify and extend.

Here we describe MOAflow, that is a bioinformatic pipeline for the analysis of MNase-defined cistrome-Occupancy (MOA-seq) data, developed on the previous work of *Liang et al., 2022*. The MOA-seq methodology was developed as a high-resolution, high-throughput, genome-wide method specifically designed to accurately identify putative Transcription Factor (TF) binding sites, called MOA-footprints (MFs), and to reveal Accessible Chromatin Regions (ACRs) in plants genomes (*Savadel et al., 2021*). MOA-seq can generate an exceptionally high volume of data (*Engelhorn et al., 2025*), especially when this technique is applied to species with very large genomes, making downstream analysis computationally demanding and complex particularly if based on separate scripts and standalone software. To overcome these constraints, the original pipeline has been re-designed using Nextflow, resulting in a modular, automated, and portable solution suitable for large-scale analyses. This new design ensures efficient resource allocation, faster execution, and reproducibility, while supporting collaborative use and workflow sharing. MOAflow was tested and benchmarked with the dataset of *Liang et al., 2022*, both in a local server and in a cloud-based environment (Microsoft Azure).

## Materials and methods

### Workffow overview

MOAflow as the original pipeline of Liang et al 2022 is built upon a wide range of *state-of-the-art* software and data resources. The pipeline consists of three main phases: data pre-processing, reads alignment, and peak calling (see **Fig. A**). During the pre-processing step, quality control (QC) is performed on the raw PE reads (Paired Ends library is mandatory as input). FastQC (*Andrews et al., 2010*) is used to check quality before and after the trimming process, which is performed by SeqPurge (*Sturm 2025*). Subsequently, overlapping PE reads are merged into single reads (SR) using FLASH (*Magoč et al., 2011*) in order to improve quality and increase the fragment length. The alignment process begins with the generation of the genome index (if necessary) using STAR (*Dobin et al., 2013*). Annotation file (gtf format mandatory) can be provided. Single End reads (SE) are aligned to the genome using STAR software. Subsequently, SAMtools (*Danecek et al., 2021*) is used to filter the generated bam files, retaining reads that have a quality score (MAPQ) of 255 and are shorter than 80 bp. The effective genome length is calculated with the script unique-kmers.py from Khmers (*Crusoe et al., 2015*), using either a fixed k-mer size or the average mapped fragment length (AMFL) calculated with SAMtools stat. bamCoverage function of deepTools (*Ramírez et al., 2016*) is used to generate normalized bedgraph files (reads per genomic coverage, RPGC) of the complete read data. Finally, the MOA footprints (MFs) discovery process involves reducing reads to 20 bp (optional) and peak calling. To generate high-resolution maps, each read can be reduced to 20 bases centered on the midpoint. The shortening is performed using a simple awk command: in the case of reads with an odd number of bases, the central base is taken, and the sequence is extended by 10 bases on each side; in the case of reads with an even number of bases, one of the two central bases is chosen at random, and the sequence is extended by 10 bases on each side. Subsequently, the reads, shortened or kept at their original length, are used by MACS3 (*Zhang et al., 2008*) for peak calling (callpeak sub-command). It should be noted that the analysis can be performed either by keeping the biological replicates separate (default case) or by merging the replicates to obtain a deeper coverage. However, if the replicates are merged, samples from different experimental conditions (e.g., treatment and control) cannot be analyzed at the same time.

**Fig. A.**
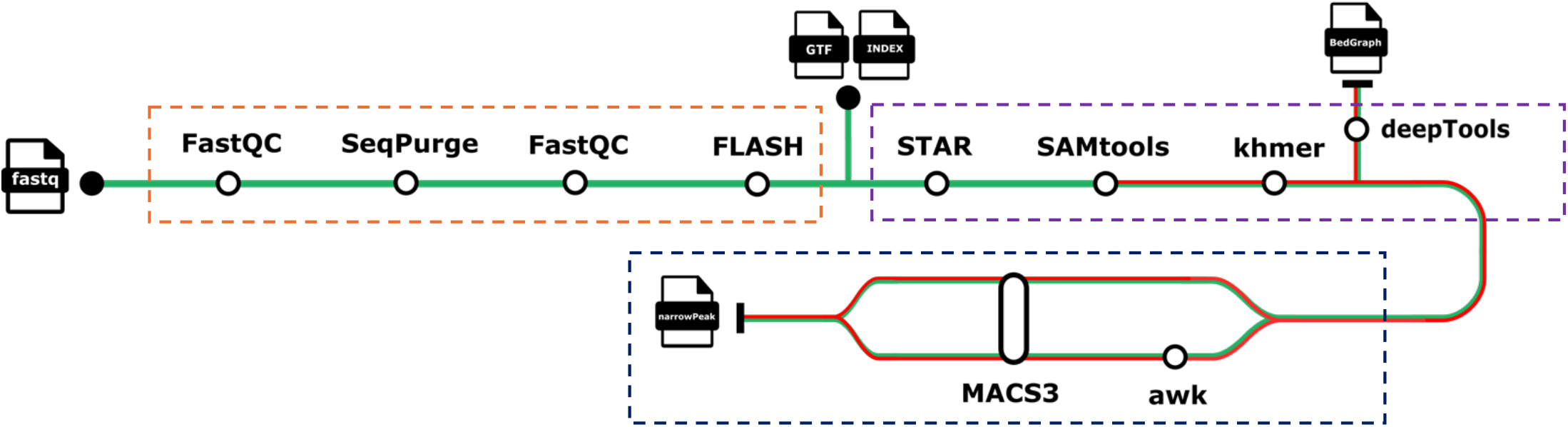
MOAflow workflow. The software used in the analysis are displayed in the metromap. The dotted boxes refer to the stages of analysis: pre-processing (orange), alignment (purple), and peak calling (blue). The green-red line starting from SAMtools means that replicates can be merged or not.

### Implementation in Nextffow

The MOAflow pipeline is implemented in Nextflow, making use of the new DSL2 syntax. Nextflow is an extension of Groovy, which in turn is a superset of Java and simplifies the writing of data-intensive computational pipelines. This pipeline is composed by 13 different modules, each one representing a step in the analysis. Modules are linked together in the workflow script, called *main*.*nf*. The *nextflow*.*config* file defines the pipeline resources and default parameters, while the *params*.*json* file is used to specify run-specific settings. Input samples are provided to the pipeline through a CSV file, in which the user defines each sample name along with the corresponding raw read files. To date, the software used in the MOAflow pipeline are run and containerized exclusively with Docker. The description of parameters and output files generated by MOAflow is given at the documentation page in the code repository.

### Computing environments

MOAflow pipeline was tested on both local machine and in cloud environment (Microsoft Azure). The local system was a Windows Server 2019 Standard (Fujitsu Primergy RX900 S2) equipped with eight Intel Xeon CPU (E7-8860 2.27GHz CPUs, 80 cores, 160 threads), 224 GB of RAM, and a storage system consisting of a local unit of 272 GB and an external high‐capacity storage unit of 21.26 TB. The run was performed in an Ubuntu v24.04.3 LTS virtual machine (made by Oracle VM VirtualBox v7.0) with 64 CPUs, 83.5GB of RAM and 2 TB of storage (vHD). The cloud environment was composed by a three‐node cluster equipped with Intel Xeon Platinum processors, each node providing 96 vCPUs and 384 GB of RAM and using a 2 TB premium file share mounted via NFS on the Azure VMs to store raw data, working directories, and output files. The Open OnDemand v4.0.0 interface was used to simplify the use of Nextflow in the cloud environment.

### MOAffow benchmarking

Since the main goal of this work was to redesign the pipeline of *Liang et al. (2022)* in Nextflow while keeping its original structure unchanged, performance evaluation was conducted using various benchmark metrics and comparison strategies. The most direct analyses involved comparing the results reported in the original work with those obtained using MOAflow. The simplest comparisons concerned the number of input reads used for alignment, the number of MFs identified per sample, and the median length of full-length or 20 bp shortened MOA peaks. For a more extensive analysis based on the genomic regions shared between the MOAflow results and those of the original study, two distinct metrics were calculated: the Jaccard index and the F1-score. The Jaccard index was calculated using the jaccard function of Bedtools (v2.31.1, *Quinlan et al., 2010*), using default parameters, on the different sets of genomic intervals to be compared. The calculation of Recall, Precision and F1-score (F1 = 2 × (Precision × Recall) / (Precision + Recall)) was performed using a combination of Bedtools intersect commands (with flags -f 0.5 -r -u) and awk commands (see *f1_score_calc*.*sh* script for more details).

## Results

### Pipeline execution

To test the pipeline performance in terms of biological results, MOAflow was run on the MOA-seq dataset from the work of *Liang et al. (2022)*, with the aim of reproduce their results. The input samples consisted of PE reads obtained from three biological replicates of B73 maize under control conditions and three other B73 biological replicates subjected to heat stress (4 h at 40 °C). The B73 AGPv4 assembly was employed as the maize reference genome for read mapping. The pipeline ran successfully and generated the expected output files. MOAflow was run with a single command, with fully automated installation and execution and an efficient task distribution across eleven different software tools to complete the analysis and provide quality control metrics, without any manual intervention required. The evaluation of pipeline performance was based on the following metrics: the number of inputs reads (with the percentage of uniquely mapped reads); the number of MOA peaks identified using full-length reads and reads trimmed to 20 bp (these called MOA footprints); and the median MOA peak length for both full-length and shortened reads. **Table X** summarizes the input read alignment results, benchmarked against the findings of *Liang et al. (2022)*. The results obtained with MOAflow, although not identical, closely correspond to those reported in the original study. The deviation in input reads per replicate ranged from 0.09% to a maximum of 0.2% fewer reads, and the unique alignment rates were also comparable. **Table Y** shows the number of MFs per sample. The results were highly consistent with the original ones, with differences in peak counts ranging from 0.02% to 0.065% more MFs. Median MOA peak lengths were consistent: 180 bp for full-length peaks versus 179 bp reported in the article and 34 bp for MFs (shortened peaks), matching the original value.

**Table X.** Number of input merged reads per replicate (unique alignment rate).

| Table X | *Liang et al., 2022* | MOAflow |
| --- | --- | --- |
| **CONTROL1** | 126.939.893 (45%) | 126.716.389 (46%) |
| **CONTROL2** | 166.432.747 (44%) | 166.101.382 (45%) |
| **CONTROL3** | 138.693.816 (40%) | 138.569.295 (45%) |
| **STRESS1** | 158.734.914 (45%) | 158.500.699 (49%) |
| **STRESS2** | 131.979.910 (46%) | 131.850.844 (50%) |
| **STRESS3** | 160.081.276 (44%) | 159.861.675 (47%) |

**Table Y.**
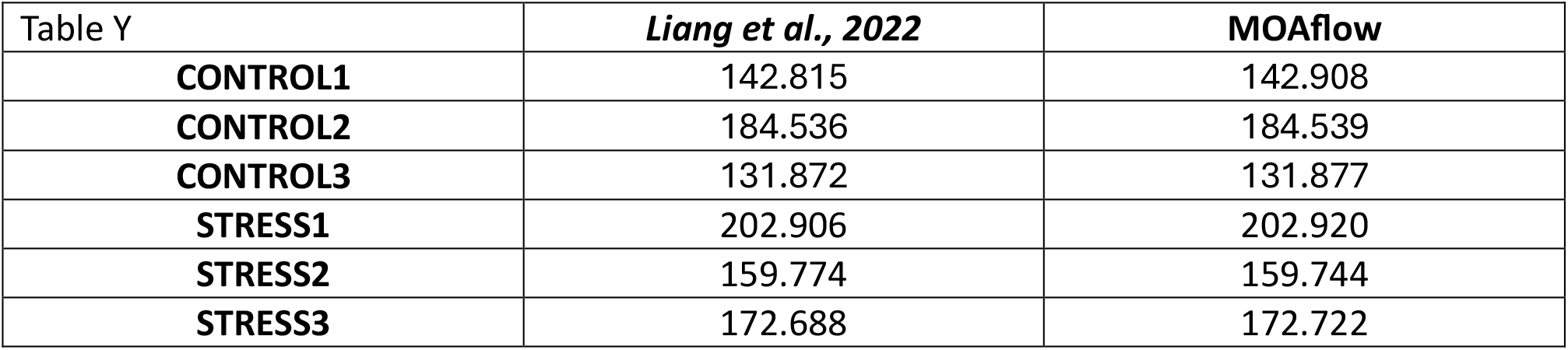
Number of MOA footprints (MFs, shortened peaks).

Although these results are consistent from a merely numerical point of view, it was necessary to assess whether the MFs found by MOAflow overlapped the same genomic regions identified by the original work. Due to the lack of MFs coordinates files, we used the MOA peaks files (generated by full length reads) from the UCSC genome browser (“Availability of data and materials” section) from *Liang et al. (2022)*. The Jaccard index, which measures the ratio between the number of intersecting base pairs between two sets and the number of base pairs in the union of the two sets, as well as the precision, recall and F1-score statistics using the results of the original study as the “gold standard”, were calculated as benchmarks. The Jaccard index, calculated with jaccard function of Bedtools (v 2.31.1), for each replicate in both conditions range from the lowest value of 0.92 (control 1) to the highest one of 0.99 (stress 3). **Table O** depicts the results of the F1-score statistic and related metrics. Subsequently, the results of the differential MOA footprints analysis between control and heat conditions (Additional file 2, Data S1: MOA‐seq footprints of B73 detected under control or heat conditions) from *Liang et al. (2022)* were used to assess the MFs resulted by MOAflow pipeline. The differential analysis was performed with the DiffBind (v2.12.0, *Stark et al., 2011*) function in R on the results of the MOAflow pipeline. Again, the Jaccard index and F1-score statistics using the results of the original study as the “gold standard” were calculated as benchmarks. **Table Z** summarizes differential MFs, and the distribution of those regions with increased accessibility in heat stress and those with less accessibility to heat stress compared to controls. The Jaccard index was calculated both on the overall set of differential MFs and separately for each experimental condition. For the total dataset, the Jaccard index was 0.89, while under heat stress and control conditions it was 0.88 and 0.90, respectively. The other metrics were calculated in the same manner and **Table J** summarizes the values. The high values of the Jaccard index as well as those of the F1-score indicate that the workflow performed as expected and it is consistent with the results of *Liang et al. (2022)*.

**Table O.** F1-score statistics related to the MOA full length peaks per replicate. The MOA full length peaks of the original article were used as gold standard, while the MOAflow results as query.

| Table O | Recall | Precision | F1-score |
| --- | --- | --- | --- |
| **CONTROL1** | 0.93 | 0.94 | 0.94 |
| **CONTROL2** | 0.99 | 0.99 | 0.99 |
| **CONTROL3** | 0.97 | 0.96 | 0.97 |
| **STRESS1** | 0.97 | 0.96 | 0.96 |
| **STRESS2** | 0.97 | 0.96 | 0.97 |
| **STRESS3** | 0.99 | 0.99 | 0.99 |

**Table Z.**
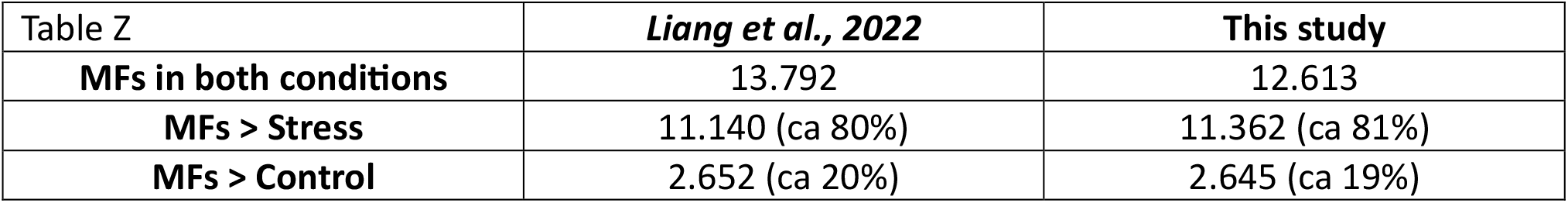
Overall of differential MFs. Distribution of those regions in increased accessibility in heat stress and those with less accessibility to heat stress compared to controls.

| Table Z | *Liang et al., 2022* | **This study** |
| --- | --- | --- |
| **MFs in both conditions** | 13.792 | 12.613 |
| **MFs > Stress** | 11.140 (ca 80%) | 11.362 (ca 81%) |
| **MFs > Control** | 2.652 (ca 20%) | 2.645 (ca 19%) |

**Table J.**
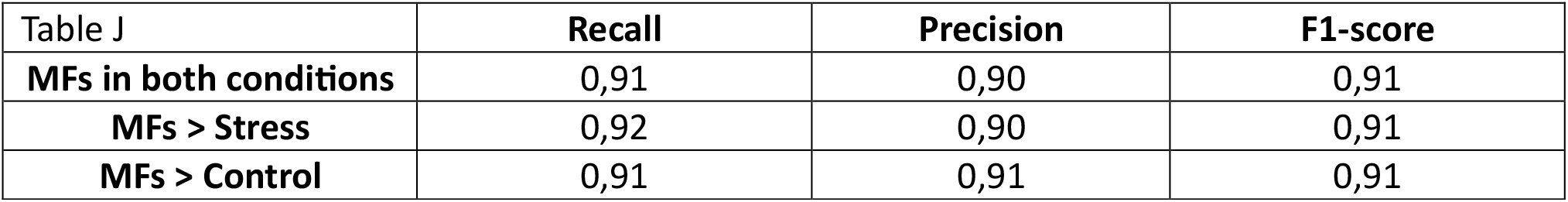
F1-score statistics calculated on differential MFs.

### Computing environments comparison

The MOAflow pipeline was evaluated in two distinct computing environments: a local server and on a cloud‐based platform (Microsoft Azure). The pipeline processed approximately 90 GB of input data, comprising raw sequencing reads and the maize reference genome, and generated roughly 229 GB of output data, including both intermediate and final files. Identical parameters, input datasets, and the execution command were used in both environments. On the local server, the run required 2 days and 4 hours, whereas the cloud ran successfully in 2 hours and 44 minutes. Total CPU usage amounted to 2,374.2 CPU‐hours on the local machine, compared with 423.4 vCPU‐hours in the cloud. In both cases, the number of tasks executed was 74.

## Discussion

As shown in the previous section, the results obtained with MOAflow are highly consistent with, and in sometimes identical to, those reported in the original study. Minor discrepancies may essentially depend on stochastic initialization, meaning that results may not be identical. This is reflected in the Jaccard index and F1-score values when comparing full-length MOA peaks and MF differential analyses. The MOAflow pipeline was evaluated across heterogeneous computing environments, revealing significant differences in execution times. The comparison aimed to assess the robustness of the pipeline in different computing context and its ability to produce comparable, and in some cases identical, results, rather than to measure performance in terms of analysis speed. The results obtained confirm this behavior.

It should be emphasized that MOAflow was intentionally not implemented with integrated differential analysis (DiffBind R package) and motif discovery, because these analyses can be highly customizable and their underlying statistical approaches require careful evaluation of the outcomes; therefore, it is preferable to perform these steps manually. Nevertheless, the MOAflow pipeline is accompanied by the DiffBind script used to validate the results. This material is provided solely as a starting point for differential analysis; consequently, it is strongly recommend adapting them to everyone specific needs.

## Data availability

The data used to test the pipeline are those of the *Liang et al. (2022)* article.

## Software availability

Source code available at: https://github.com/Jacopot12/MOAflow. License: MIT License.

## Declaration of AI use

Copilot was used to improve the readability and language of the manuscript. The authors take full responsibility for the work.

## References

Simon Andrews, F. K., et al. “FastQC: a quality control tool for high throughput sequence data.” 2010.

Berger, Bonnie, and Yun William Yu. “Navigating bottlenecks and trade-offs in genomic data analysis.” Nature Reviews Genetics 24.4 (2023): 235–250.

Crusoe et al., The khmer software package: enabling efficient nucleotide sequence analysis. 2015. 10.12688/f1000research.6924.1

Danecek, Petr, et al. “Twelve years of SAMtools and BCFtools.” Gigascience 10.2 (2021): giab008

Di Tommaso P, Chatzou M, Floden EW, et al.: Nextflow enables reproducible computational workflows. Nat Biotechnol. 2017; 35(4): 316–319.

Dobin A, et al. STAR: ultrafast universal RNA-seq aligner. Bioinformatics. 2013;29:15–21.

Engelhorn, Julia, et al. “Genetic variation at transcription factor binding sites largely explains phenotypic heritability in maize.” Nature Genetics (2025): 1–10.

Jones, D. Marc, and Klaas Vandepoele. “Identification and evolution of gene regulatory networks: insights from comparative studies in plants.” Current opinion in plant biology 54 (2020): 42–48.

Liang, Zhikai, et al. “Mapping responsive genomic elements to heat stress in a maize diversity panel.” Genome biology 23.1 (2022): 234.

Magoc T, Salzberg SL. FLASH: fast length adjustment of short reads to improve genome assemblies. Bioinformatics. 2011;27:2957–63

Quinlan Aaron R., and Ira M. Hall. “BEDTools: a flexible suite of utilities for comparing genomic features.” Bioinformatics 26.6 (2010): 841–842.

Ramírez, Fidel, Devon P. Ryan, Björn Grüning, Vivek Bhardwaj, Fabian Kilpert, Andreas S. Richter, Steffen Heyne, Friederike Dündar, and Thomas Manke. deepTools2: A next Generation Web Server for Deep-Sequencing Data Analysis. Nucleic Acids Research (2016). doi:10.1093/nar/gkw257.

Savadel, Savannah D., et al. “The native cistrome and sequence motif families of the maize ear.” PLoS genetics 17.8 (2021): e1009689.

Stark R, Brown G (2011). DiffBind: differential binding analysis of ChIP-Seq peak data. http://bioconductor.org/packages/release/bioc/vignettes/DiffBind/inst/doc/DiffBind.pdf.

Marc Sturm (2025) «imgag/ngs-bits: 2025_03». Zenodo. doi: 10.5281/zenodo.15051584.

Zhang, Yong, et al. “Model-based analysis of ChIP-Seq (MACS).” Genome biology 9.9 (2008): R137.

